# An Unexpected Single-step Glycosylation Enables the Construction of Antibody-Biomacromolecule Conjugates as Therapeutics

**DOI:** 10.1101/2022.09.04.506510

**Authors:** Yang Yang, Zhentao Song, Tian Tian, Zihan Zhao, Ji Chen, Jiangping Hu, Xin Jiang, Guoli Yang, Qi Xue, Xinlu Zhao, Wanxing Sha, Yi Yang, Jie P. Li

**Author notes:** Contributed equally (Yang Yang, Zhentao Song, Tian Tian).

## Abstract

Glycoengineering has been demonstrated to be a powerful tool to construct site-specific antibody conjugates. However, with most glycoengineering strategies, several hours to days are needed to complete the reaction, and the payloads are limited to small molecules. Herein, we show that reducing the complexity of Fc glycan could dramatically boost the enzymatic glycoengineering of IgG molecules to achieve unexpectedly high efficiency and payload capacity. Notably, antibodies with the modified Fc glycan were successfully conjugated with biomacromolecules, *e.g*., oligonucleotides and nanobodies, in one step by an engineered fucosyltransferase. Moreover, the conversion of this chemoenzymatic engineering could be finished in minutes, or up to hours, affording homogeneous products. The trisaccharide linker of these antibody conjugates exhibits excellent hydrophilicity and stability and reduced ADCC activity. Using this platform, we successfully synthesized an antibody-conjugate-based HER2/CD3 bispecific antibody and applied it to selectively destroy patient-derived cancer organoids by reactivating endogenous T cells inside the organoid. These results suggest that this bispecific antibody format has a high translational value.

---

Antibody conjugates are the major forms of antibody-derived macromolecules for the targeted manipulation of biological systems, which combines the exquisite specificity of antibodies with the distinct functions of payloads via chemical conjugations. In recent years, antibody‒drug conjugates (ADCs) have been very successful in cancer treatment^1^, especially trastuzumab deruxtecan (T-DXd; DS-8201), an ADC composed of an anti-HER2 antibody and topoisomerase I inhibitor, which might redefine the standard therapy of HER2-low breast cancer^2^. Furthermore, the first antibody– oligonucleotide conjugate (AOC) entered into a phase I clinical trial last year, and this breakthrough broadens the molecular types of antibody conjugate-based drugs^3^. Beyond therapeutic application, AOCs have been used in the sensitive and multiplexed detection of proteins in single cells^4^ or traceless samples^5^; for example, AOCs are used to rapidly measure protein biomarkers in a minimal sample volume from patients^6^. It is likely that antibody conjugates will be extensively expanded in the near future for a broad range of applications, including applications with small-molecule drugs, fluorophores, catalysts, synthetic polymers, oligonucleotides, peptides, enzymes and other proteins, either as technical tools or clinical therapeutics^7–11^. Thus, the future requirements of antibody conjugation involve satisfying the need for a versatile and efficient platform in which homogenous antibody conjugates with unlimited payloads can be constructed^12^.

The N-linked glycan on asparagine 297 (N297) in the Fc domain was found to be a single and universally conserved glycosylation site of IgG molecules. This glycosylation is predominantly composed of varied amounts of N-acetylglucosamine, fucose, galactose, mannose and N-acetylneuraminic acid (sialic acid) residues, which are assembled in different complex-type biantennary structures^13^. The inherent site specificity of Fc glycosylation makes glycoengineering a universal and efficient approach to synthesizing homogeneous antibody conjugates^10, 14^. Notably, due to the great translational potential, several antibody conjugates generated by glycoengineering have been used in clinical trials by biotechnology companies. In general, current platforms mainly utilize galactosyltransferases (GalTs)^15, 16^, sialyltransferases (STs)^17^ and endoglycosynthases (EndoGSs)^18–21^ derived from endoglycosidases to chemoenzymatically synthesize antibody conjugates via Fc glycan remodeling. Installing the payload via these platforms usually follows a two-step procedure^10^, in which bioorthogonal reaction handles, such as azides, were first introduced onto Fc via unnatural glycosylation, and drugs were further installed by bioorthogonal reactions; in addition, one-step EndoGS-based methods were recently reported^22–24^. However, with most of these platforms, several hours to days are needed to transfer small-molecule payloads, and the platforms are limited to the direct transfer of unnatural donor substrates that contain biomacromolecules (>10,000 Da) in one step.

Fucose usually appears as the capping sugar in natural polysaccharides. The fucosylation reaction, in which a fucose could be transferred to the N-acetylgalactosamine (LacNAc, acceptor) from guanosine diphosphate β-L-fucose (GDP-Fuc, donor) via fucosyltransferase (FT), has not been extensively characterized in the context of antibody conjugation, in which fucose derivatives are transferred to the Fc glycan. We have previously reported several cases of unnatural fucosylation to engineer cell-surface LacNAc with large payloads in minutes^25^. However, directly using this reaction system for synthesizing antibody glyco-conjugates involves a relatively low efficiency, which achieves only ~40% conversion even after 16 hours (**Figure S1-2**). Herein, we demonstrated that reducing the complexity of Fc glycan by enzymatic reactions dramatically accelerated unnatural Fc fucosylation to install diversified functionalities (**Figure 1A**). Compared to the complex natural biantennary N-glycan, the simple monoantennary disaccharide LacNAc on N297 of Fc could be modified via an optimized 1,3-FT-mediated fucosyl-biotinylation with a much higher reaction rate (minutes versus hours). More importantly, biomacromolecules, such as proteins (~15 kDa) and oligonucleotides (~100 nt), could be transferred to this created simple LacNAc unit on IgG in one step. Further successful applications of these conjugates demonstrated the general applicability of this platform and provided distinct molecular entities for future therapeutics. In particular, a HER2/CD3 bispecific antibody was successfully synthesized in this platform, which enables the recruitment of endogenous cytolytic T cells to kill tumor cells in human tumor organoids (**Figure 1B**).

**Figure 1.**
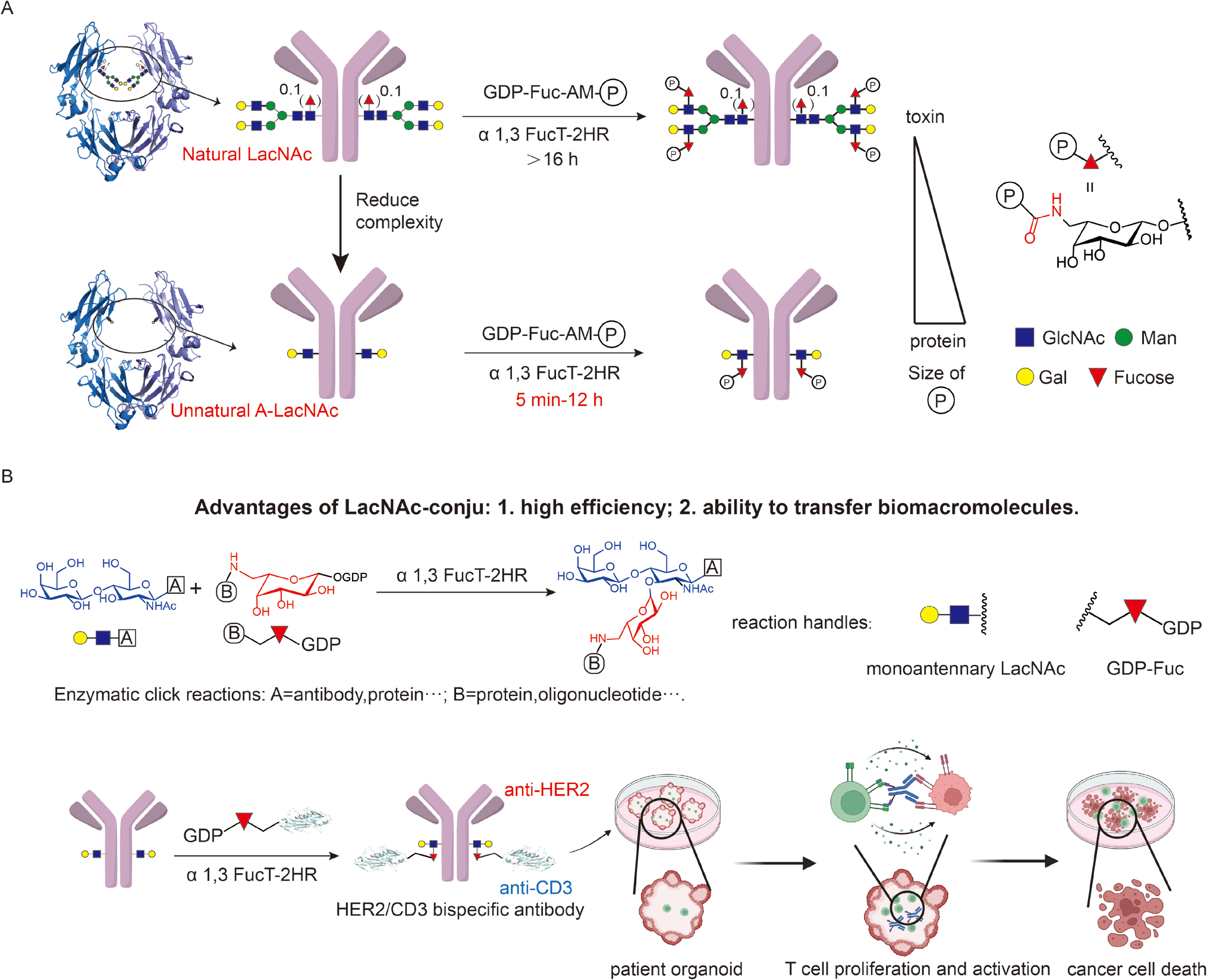
Schematic illustration of the LacNAc-conju platform for antibody-biomacromolecule conjugate construction. (A) Schematic representation of enzymatic unnatural A-LacNAc labeling (LacNAc-conju) for constructing antibody conjugates. By reducing the Fc glycan complexity, biomacromolecules are transferred to antibodies with high efficiency. (B) Chemical synthesis of HER2/CD3 bispecific antibody via the LacNAc-conju platform provides a novel chemical bispecific antibody that enables the recruitment of cytolytic T cells for tumor killing.

## Results and discussion

### Optimization of FT-mediated LacNAc labeling on the natural Fc glycan of the antibody

Recently, an FT enzyme originating from *H. pylori 26695* was reported to label the LacNAc unit efficiently on cell-surface glycoproteins or glycolipids with functional moieties^25^. To utilize this reaction for the modification of Fc glycan on an antibody, we chose an anti-human epidermal growth factor receptor 2 (HER2) antibody, trastuzumab (Tras), with engineered homogeneous glycan structure (G2F, 4 LacNAc per antibody) as the model protein (**Figure S1**). Using our previously reported FT enzyme and GDP-Fuc-Azide-derived biotin probe (triazole linker, TA), LacNAc on the G2F-Tras antibody were only partially fucosylated (~1.4 biotin per antibody) under saturated reaction conditions (5 mM GDP-Fuc-TA-Biotin and 16 hours) (**Figure S2**). To achieve high efficiency of FT-mediated LacNAc labeling, the linker between GDP-Fuc and the payload as well as the sequence of the FT enzyme were carefully optimized^26, 27^. Notably, the use of GDP-Fuc-Biotin bearing an amide linker (GDP-Fuc-AM-Biotin) and an FT variant with shorter heptad repeats (FT-2HR) showed significantly improved reaction efficiency (**Figure S2C, S3**). As a result, we could modify G2F-Tras antibodies with four almost exclusive biotin probes via fucosylation using the improved substrate and enzyme (**Figure S3**).

### Reducing Fc glycan complexity boosts enzymatic A-LacNAc labeling (LacNAc-conju)

Although we achieved one-step enzymatic labeling of Fc glycan, the reaction was still inefficient and needed 16 hours to complete (**Figure 2A**). In contrast, transferring GDP-Fuc derivatives to LacNAc on the cell surface by FT required only 10 minutes^25^. According to the crystal structure of the antibody Fc domain, the glycans are positioned between two CH2/CH3 domains and fill the large pocket between the two CH2 domains^28^ (**Figure 2B**). We envisioned that the extent of crowding in the pocket filled by the double biantennary glycans hindered LacNAc enzymatic labeling. Therefore, we reduced the complexity of the Fc glycan using EndoS (an endoglycosidase used for the selective hydrolysis of the glycans at N297 in antibodies) and AlfC (an α-fucosidase to trim α(1,6)-fucose) to create the GlcNAc monosaccharide on N297^29, 30^, which occurs due to the trimming of biantennary glycan and core fucose. After that, we used bovine β-1,4-galactosyltransferase 1 (Y289L) (β_4_GalT1) to perform galactosylation with the exposed GlcNAc and created a monoantennary disaccharide structure, LacNAc, on N297 of the Fc domain (hereafter named A-LacNAc because it directly links the amino acid) (**Figure 2B**). Although three enzymes are needed for the conversion, the trimming and transferring processes could be realized in one pot by the addition of EndoS, AlfC, β_4_GalT1 and uridine diphosphate galactose (UDP-galactose) together. After the one-pot reaction, the resulting antibodies with A-LacNAc (Tras-(A-LacNAc)_2_) were purified by Protein A resin and further modified with biotin moieties by GDP-Fuc-AM-Biotin (80 μM). To our surprise, the transfer of fucose-biotin to A-LacNAc reached almost 100% conversion within only 5 minutes (**Figure 2B**), which outperformed those with complex biantennary structures (16 hours were needed even with 5 mM GDP-Fuc-AM-Biotin) (**Figure 2A**). Finally, we monitored the conversion efficiency of natural LacNAc and A-LacNAc at different intervals with the same substrate concentration (**Figure 2C-D**). More than 70% A-LacNAc was converted within only 0.5 minutes and was almost completely converted within 5 minutes, while only ~20% natural LacNAc was converted even after 5 hours (**Figure 2C-D**). The significant enhancement in reaction efficiency indicates that the crowded structure of natural Fc glycan is the main barrier for glycoengineering, and reducing Fc glycan complexity could greatly boost the enzymatic reaction. Thus, we named this platform “LacNAc-conju” since it enables the modification of A-LacNAc on the antibody Fc domain.

**Figure 2.**
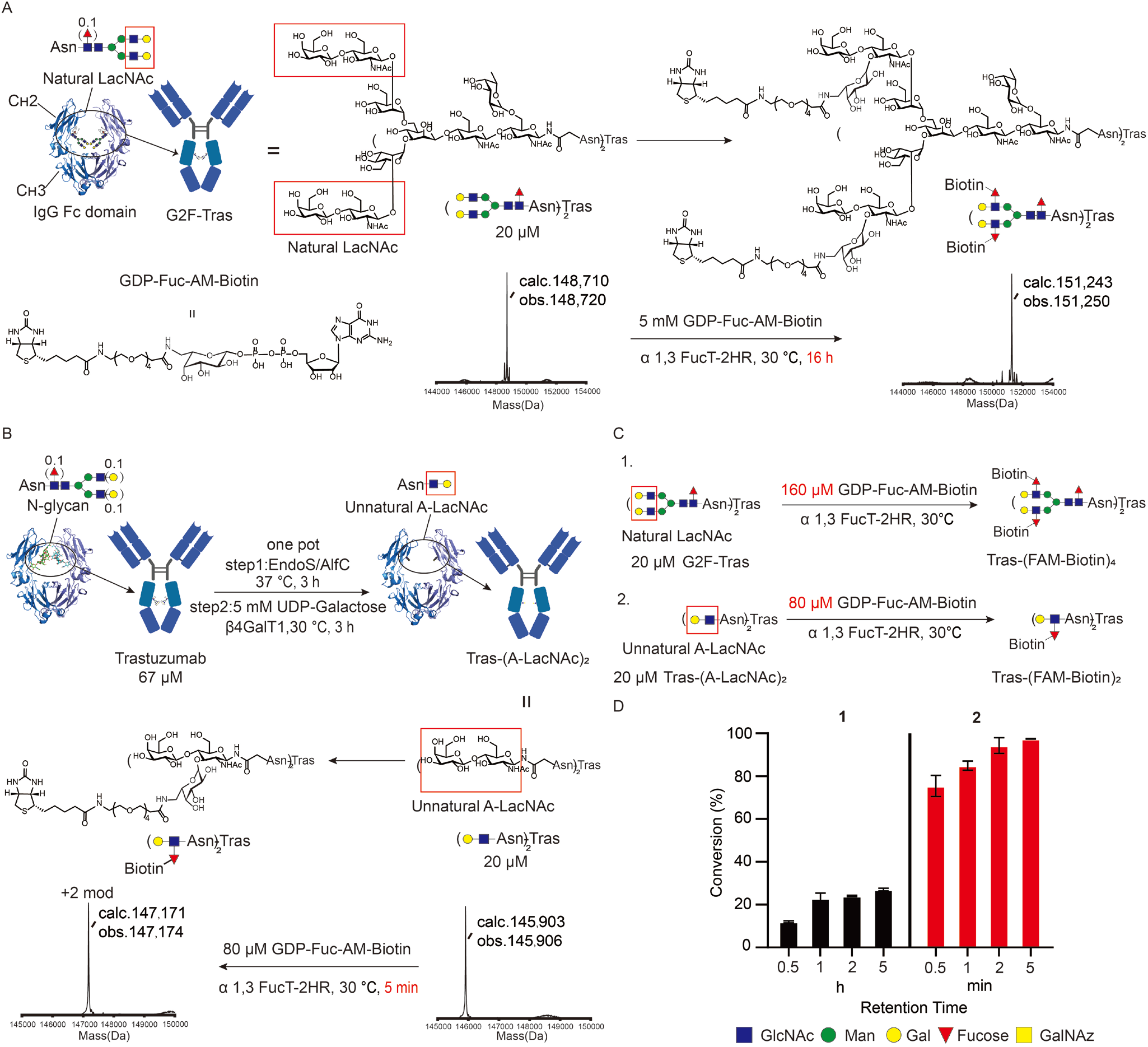
Reducing the Fc glycan complexity greatly improves the efficiency of antibody glycoengineering. (A) For antibody biotinylation on the natural biantennary glycan structure (four LacNAc units per antibody) mediated by α 1,3 FucT-2HR, 16 hours were needed to complete the conversion. obs: Observed; calc: Calculated. (B) Biantennary glycan on antibody was reengineered to generate a simple monoantennary disaccharide on N297 by cascade enzymatic reactions. Due to the reduced complexity of Fc glycan, A-LacNAc is a more favorable substrate for α 1,3 FucT-2HR. Only 5 minutes were necessary for full antibody biotinylation on A-LacNAc, and much less donor substrate was needed. This reaction cascade could be combined in one pot. (C-D) The time-dependent conversion efficiency to obtain Tras-(FAM-Biotin)_4_ (reaction 1) and Tras-(FAM-Biotin)_2_ (reaction 2). (C) Scheme of the one-step synthesis of different biotinylated trastuzumab derivatives. Reactions (1 and 2) were performed in the presence of 20 μM substrate (G2F-Tras, Tras-(A-LacNAc)_2_) and 0.8 mg/mL enzyme in Tris-HCl buffer (25 mM, pH 7.5) at 30 °C. (D) The time-dependent conversion efficiency of reactions 1 and 2. Reactions were monitored by LC‒MS.

To further compare the reaction rate of LacNAc-conju with the widely used click chemistry, *e.g*., strain-promoted azide-alkyne cycloaddition (SPAAC), we synthesized the Tras antibody, which was modified by a simple unnatural disaccharide that contained the azide group, as the model substrate to react with a dibenzocyclooctyne (DBCO)-biotin probe. The results suggest that LacNAc-conju exhibits higher efficiency than that of SPAAC (**Figure S4**), demonstrating the power of unnatural enzymatic reactions for the highly efficient modification of proteins. More broadly, the LacNAc-conju platform was not restricted to biotin probes and could be generally applicable to different IgG-based drugs, e.g., durvalumab (**Figure S5-6**).

### Characterizing the pharmaceutical property of antibody conjugates with a trisaccharide linker

Although antibody conjugates are efficiently constructed with LacNAc-conju, the created trisaccharide linker is a novel glycan structure (Lewis X linked to the amino acid directly, named as A-LeX) and needs to be characterized before further application. To test its property, we used ADC as an example of different conjugates. First, monomethyl auristatin E (MMAE), conjugated to the most common cleavable linker, Val-cit-PAB (Vc-PAB), was synthesized and modified with GDP-Fuc as described in the methods (**Figure 3A**). As expected, the generated GDP-Fuc derivative (GDP-Fuc-AM-VcPAB-MMAE) showed excellent hydrophilicity and could be used to synthesize a homogeneous ADC, Tras-(A-LeX-MMAE)_2_, from Tras-(A-LacNAc)_2_ (**Figure 3A, S7A**). In fact, the hydrophobicity of MMAE has been reported to induce the quick plasma clearance and then could reduce exposure and *in vivo* efficacy^31^. To our delight, further analysis of Tras-(A-LeX-MMAE)_2_ by hydrophobic interaction chromatography (HIC) and size exclusion chromatography (SEC) demonstrated that the Tras antibody was uniformly labeled with two MMAE toxins, and no aggregation was observed (**Figure S7B-C**). Moreover, Tras-(A-LeX-MMAE)_2_ with a trisaccharide-linker showed much improved hydrophilicity compared to that of the maleimide-based linker, which was even very close to that of the unmodified antibody (**Figure S7B**). After obtaining this ADC, we next characterized the stability of this trisaccharide linker via enzymatic trimming and found that neither PNGase F nor PNGase A cleaved this trisaccharide on Fc (**Figure 3B, S7D**). This result suggests that antibody conjugates from the LacNAc-conju platform exhibit potential resistance toward glycosidases. Additionally, Tras-(A-LeX-MMAE)_2_ was highly stable in human serum within 8 days (**Figure 3C**), indicating that this glycan linker is also resistant to other enzymes and nucleophiles in serum. Furthermore, it is well established that performing Fc glycosylation with an antibody significantly affects its affinity for FcγRIIIa receptors and antibody-dependent cellular cytotoxicity (ADCC). Thus, we characterized the ADCC effect of the trisaccharide-linked ADC, and it displayed a much lower ADCC effect due to the loss of the biantennary structure, which is similar to the Tras antibody that contains the A-LacNAc Fc glycosylation (**Figure 3D, S7E**). Finally, the function of Tras-(A-LeX-MMAE)_2_ was also confirmed *in vitro* and *in vivo*; this antibody maintained the same binding capacity as the parent antibody and showed superior antitumor activity (**Figure 3E, S7F-G**). Taken together, the results indicated that by using ADCs as the drug model, the good pharmaceutical properties of antibody conjugates bearing the trisaccharide linker was demonstrated.

**Figure 3.**
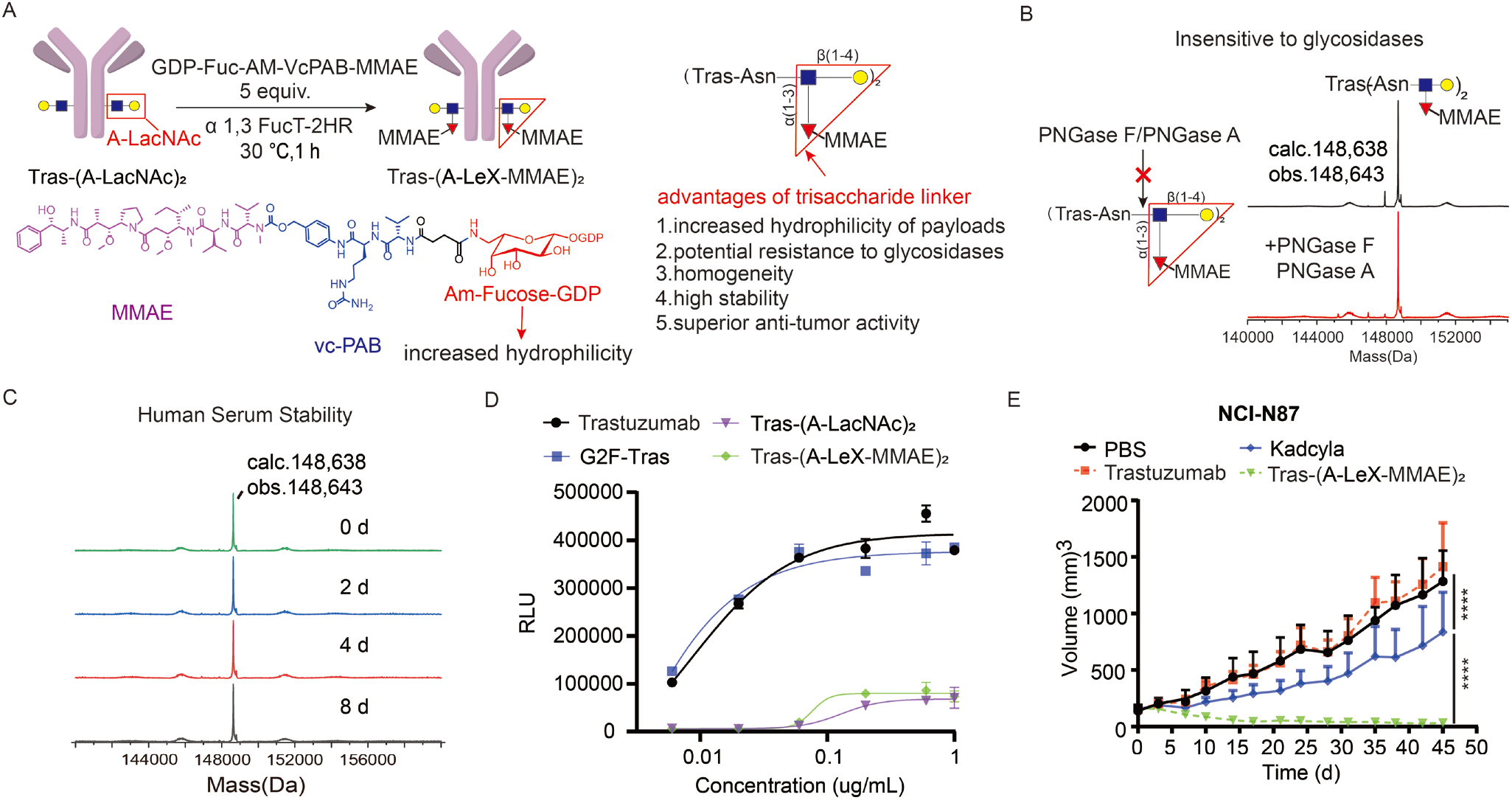
Characterization of the trisaccharide linker of LacNAc-conju. (A) Scheme of the one-step synthesis of the site-specific ADC, Tras-(A-LeX-MMAE)_2_, using LacNAc-conju. (B) ESI-MS analysis of Tras-(A-LeX-MMAE)_2_ before and after PNGase F and PNGase A treatment. (C) The stability of Tras-(A-LeX-MMAE)_2_ in serum characterized by ESI-MS analysis after 0, 2, 4, 6, and 8 days of incubation. Deconvoluted spectra are shown. obs: Observed; calc: Calculated. (D) Antibody-dependent cellular cytotoxicity (ADCC) of different trastuzumab glycoforms. ADCC activity was assessed by a robust ADCC assay using a reporter Jurkat cell line (expression of human FcγRIIIa, high affinity (V158) variant and Fcγ chainas). RLU: relative luciferase units. (E) *In vivo* antitumor activity of Tras-(A-LeX-MMAE)_2_ in a nude mouse human gastric NCI-N87 xenograft model. Mice were *i.v*. injected with PBS, trastuzumab (5 mg/kg), Kadcyla (5 mg/kg) or Tras-(A-LeX-MMAE)_2_ (5 mg/kg) when the average tumor size reached 150-200 mm^3^ (n = 5 mice per group).

### Use of GDP-Fuc-ssDNA to test the payload capacity of the LacNAc-Conju platform

In addition to the unexpectedly high efficiency of the LacNAc-conju platform for modifying antibodies with small molecules, we explored the donor substrate tolerance of 1,3-FT-2TR to test the payload capacity of this platform. By increasing the molecular size of single-stranded DNA (ssDNA), we created a library of “molecular rules” ranging from 8,000 Da to 35,000 Da, and these probes contained a 5’ amine group for further modification (**Figure 4A**). After that, GDP-Fuc was introduced to these ssDNA probes (20 nt, 30 nt, 42 nt, 59 nt, 85 nt and 106 nt) by combining amine coupling and inverse-electron-demand Diels-Alder (iEDDA) reaction^32^ and confirmed by LC‒MS analysis (**Figure 4A, S8**). Then, we used these oligos to synthesize AOCs on Tras-(A-LacNAc)_2_ antibody and tried to explore the maximal length of ssDNA that could be transferred by LacNAc-conju (**Figure 4B**). As revealed by SDS‒PAGE gel analysis, Tras-(A-LacNAc)_2_ conjugation was completed with all of these oligos by using 1,3-FT-2TR, in which almost no unmodified heavy chain was observed in each lane (**Figure 4C, S9A-B**). The specificity of Fc modification and the oligo to antibody ratio (OAR) were confirmed by gel analysis and LC‒MS analysis, suggesting that only the heavy chain the antibody was modified with one ssDNA probe (**Figure 4C-D, S9C-F**). Noteworthily, we failed to characterize the molecular weight of AOC that was conjugated to a long ssDNA (>50 nt) by LC‒MS, which was because negatively charged oligos neutralized the positive charges of the antibody (**Figure S9G-J**). Moreover, the conjugate of ssDNA larger than 100 nt is also difficult to resolve by SDS‒PAGE. Thus, we could not characterize the maximal payload capacity of LacNAc-conju platform. However, we believe that a payload capacity up to 35 kDa could enable the construction of antibody conjugates with many therapeutic entities, such as siRNA, nanobody and cytokines.

**Figure 4.**
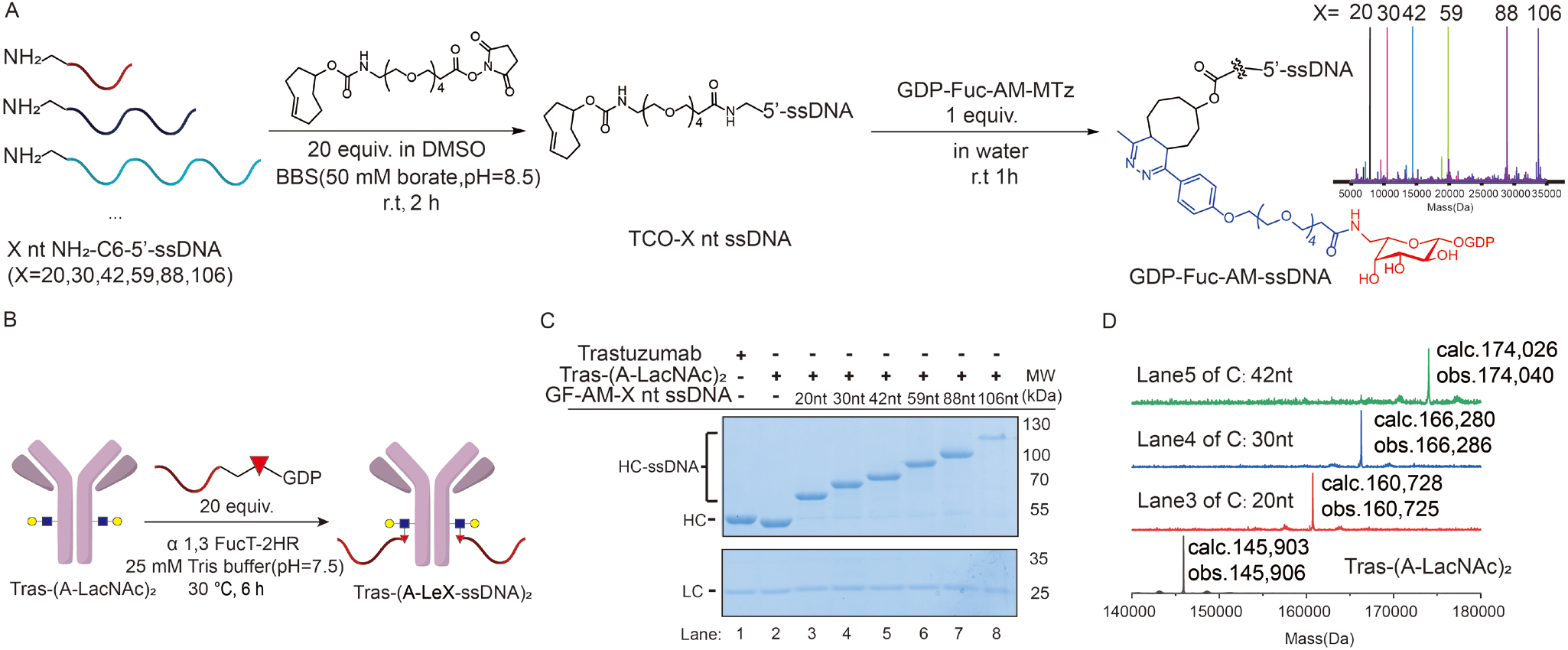
Testing the payload capacity of the LacNAc-Conju platform. (A) Chemical synthesis of GDP-Fuc-AM-ssDNA (X nt, X=20, 30, 42, 59, 88, 106). (B) Schematic representation of the synthesis of Tras-(A-LeX-ssDNA)_2_ from Tras-(A-LacNAc)_2_ antibody via LacNAc-Conju. (C) Evaluation of the conjugation efficacy by SDS‒PAGE. All the proteins were analyzed in reduced form. Lane 1: Trastuzumab. Lane 2: Tras-(A-LacNAc)2. Lanes 3-8: Crude conjugation products of Tras-(A-LeX-X nt ssDNA)_2_ (x=20, 30, 42, 59, 88, 106). HC: heavy chain, LC: light chain. (D) ESI-MS analysis (after deconvolution) of the intact purified product of Tras-(A-LeX-X nt ssDNA)_2_ made from Tras-(A-LacNAc)_2_ in one step (x=20, 30, 42).

### Construction of bispecific antibodies via LacNAc-conju

After demonstrating that the one-step transfer of biomacromolecules by FT to Fc was achieved in a site-specific and homogeneous manner by the LacNAc-conju platform, we applied the platform to construct bispecific antibodies. The bispecific T-cell engager BiTE is a type of bispecific antibody that has been leveraged to recruit cytolytic T cells for killing tumors^33^. Specifically, T cells were activated by the crosslinking of CD3 clustering on T cells, which was induced by BiTE, as well as the target antigen on tumor cells^33, 34^. As revealed by recent work, the molecular structure of bispecific or trispecific antibodies determines drug efficacy, especially for T-cell engagers^35–37^. Chemical synthesis of multispecific antibodies provides an alternative approach to designing novel molecular structures to induce better cancer-killing activity^38^. Herein, we constructed bispecific antibodies that target CD3 and HER2 antigens via the LacNAc-conju platform. First, we expressed an anti-CD3 nanobody with an LPETGG tag at the C-terminus and engineered this nanobody with DBCO via sortase A-mediated ligation^39, 40^. After that, GDP-Fuc-AM-Azide was allowed to react with DBCO-anti-CD3 nanobody, which generated anti-CD3 antibody-GDP-Fuc conjugates (GF-anti-CD3) (**Figure 5A**). Subsequently, the bispecific antibody (Tras*anti-CD3) was constructed via the LacNAc-conju reaction between GF-anti-CD3 and Tras-(A-LacNAc)_2_ (**Figure 5B**). The results of SDS‒PAGE and LC‒MS characterization showed that the reaction attained a high yield (>90%) (**Figure 5C-D**), indicating that we could realize highly efficient one-step transfer of proteins (~15 kDa) to antibodies using LacNAc-conju and provided a powerful tool for the chemical construction of bispecific antibodies.

**Figure 5.**
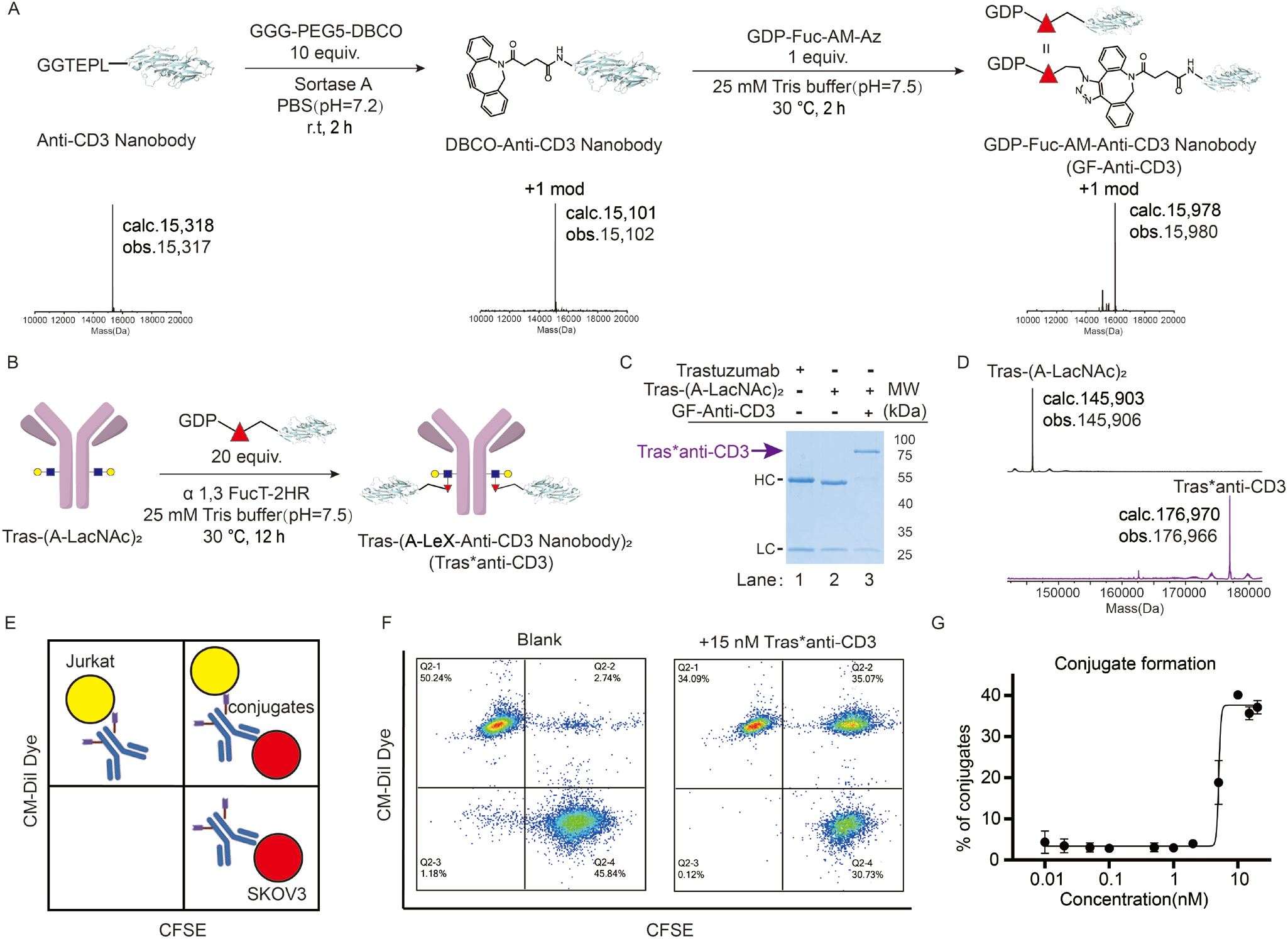
Construction and characterization of HER2/CD3 bispecific antibodies by LacNAc-conju. (A) Synthesis and ESI-MS characterization of the GDP-Fuc-AM-Anti-CD3 Nanobody. (B) Synthesis of site-specific HER2/CD3 bispecific antibody, Tras-(A-LeX-Anti-CD3 Nanobody)_2_ (Tras*anti-CD3), via LacNAc-conju. (C) SDS‒PAGE analysis of purified Tras*anti-CD3. All the proteins were analyzed in reduced form. Lane 1: Trastuzumab. Lane 2: Tras-(A-LacNAc)_2_. Lane 3: purified Tras*anti-CD3. HC: heavy chain, LC: light chain. (D) ESI-MS analysis of Tras*anti-CD3 based on Tras-(A-LacNAc)_2_. (E) Schematic illustration of Tras*anti-CD3-induced cell conjugates between Jurkat and SKOV3 cells. (F) Flow cytometric analysis of Tras*anti-CD3-induced formation of conjugates. The unconjugated cells (upper left and lower right quadrants) display mono-positive fluorescence, whereas the conjugated cells (upper right quadrant) display dual-positive fluorescence. The conjugation formation frequency (right) was measured as the fraction of cells in the upper right region. Each point represents the average frequency with two technical replicates. Data are shown as the means ± SDs. (G) Concentration-dependent cell conjugate formation induced by Tras*anti-CD3. The concentration of Tras*anti-CD3 ranged from 0.01 nM to 20 nM.

The functionality of the Tras*anti-CD3 bispecific antibody was further characterized. First, the antigen-binding capability toward HER2 and CD3 was tested by flow cytometry analysis using SKOV3 cell lines (which expressed human HER2 antigen) and Jurkat cell lines (which expressed CD3 antigen), respectively. The bispecific antibody showed a comparable binding ability to that of unmodified trastuzumab, indicating that the constructed bispecific antibody maintained the inherent binding affinity of antibodies (**Figure S10A**). For CD3, the binding ability of Tras*anti-CD3 was enhanced in comparison with that of a single anti-CD3 nanobody, which could be attributed to the multivalency of two CD3 binding sites with structural stability on Tras*anti-CD3 (**Figure S10B**). Furthermore, we characterized the bispecific antibody-mediated recruitment of T cells to cancer cells (**Figure 5E**). Jurkat and SKOV3 cells were labeled with different fluorescent cell trackers, and then 15 nM Tras*anti-CD3 was added to induce engagement. As expected, the ratio of cell conjugates was 3% without bispecific antibodies, while up to 35% cell conjugates were formed by the addition of Tras*anti-CD3 (**Figure 5F**). Moreover, the frequency of cell conjugates increased with increasing concentrations of Tras*anti-CD3, indicating that the formation of bispecific antibody-mediated cell‒cell conjugates was dose dependent (**Figure 5G**).

### Tras*anti-CD3 induces killing of cancer cells in organoids via selective T cell activation

To further assess the potency of Tras*anti-CD3 against cancer cells, we chose a model involving the coculture of primary human T cells and SKOV3 cells under antibody conjugate treatment, in which T cells were from human peripheral blood mononuclear cells (PBMCs). According to the mechanism of BiTE, Tras*anti-CD3 could induce the engagement of T cells and SKOV3 cells and subsequently trigger the activation of T cells via the formation of immune synapses (**Figure 6A**). As expected, the expression of CD69 and CD25, which are markers of T-cell activation, along with the secretion of cytotoxic cytokines (IFN-γ and TNF-α), were highly upregulated in both CD4+ and CD8+ T cells when Tras*anti-CD3 was added to the mixture of PBMCs and SKOV3 cells (**Figure 6B-C, S10D-E**). In contrast, in the absence of SKOV3 cells, Tras*anti-CD3 induced only a very weak activation of T cells, especially in the secretion of IFN-γ (**Figure 6B-C**). For comparison, Tras, anti-CD3 nanobodies and their mixtures were added to the coculture system, and none of them exhibited an ability to activate T cells. The results confirm that T activation and cytotoxic cytokine secretion in this system depend on the Tras*anti-CD3-mediated engagement of T cells and cancer cells, which is highly specific. Afterward, we used a real-time cell analyzer (RTCA) to monitor cancer cell killing induced by cytotoxic T-cell activation. To our delight, Tras*anti-CD3 displayed a dramatic potency of killing tumor cells, which outperformed Tras as well as the combination of Tras and anti-CD3 (**Figure 6D**); this activity occurred because Tras*anti-CD3 utilizes the mechanism of cytotoxic T-cell activation, while Tras depends on the ADCC effect via natural killer (NK) cells. Notably, the trisaccharide linker of Tras*anti-CD3 has been demonstrated to almost block ADCC activity, as shown in **Figure 4D** and **Figure S7E**, indicating that the potent killing of Tras*anti-CD3 treatment is only dependent on T cells. For comparison, MDA-MB-468 cells (HER2 negative) were treated under the same conditions. As a result, only nonspecific killing induced by PBMCs could be observed in each group (**Figure 6E**), which further confirms the antigen-dependent killing in this system.

**Figure 6.**
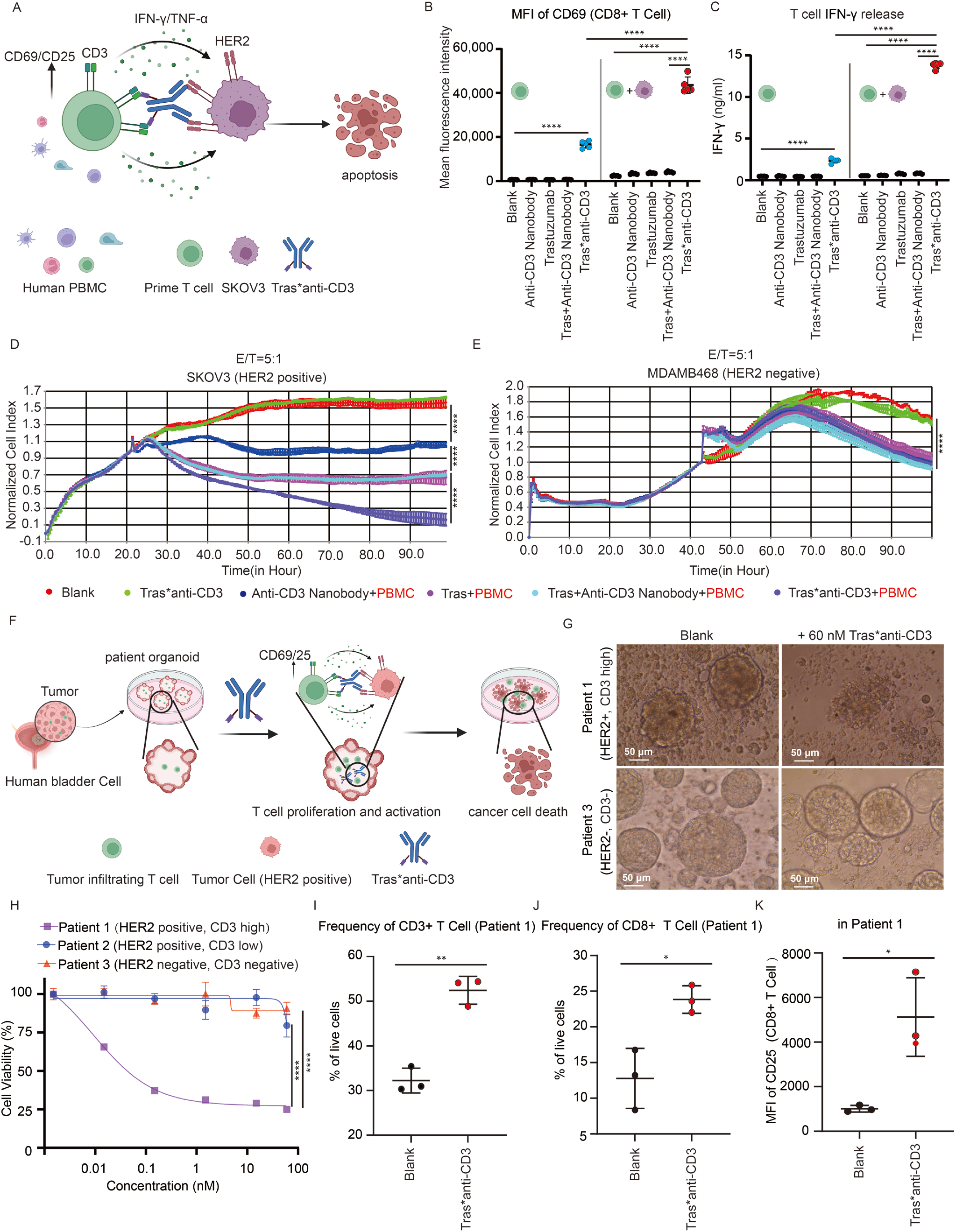
Specific immune activation and cancer killing mediated by Tras*anti-CD3 as a T-cell engager. (A) Schematic representation of cytolytic T cells and cancer cell engagement mediated by Tras*anti-CD3. (B-C) Flow cytometric analysis of CD69 expression on CD8^+^ T cells and (C) their IFN-γ secretion after 48-hour treatments (Student’s *t* test, ****P<0.0001, n=5). SKOV3 cells were added for coculturing as indicated. (D-E) Real-time cytotoxicity analysis of Tras*anti-CD3 toward SKOV3 cells (D) and MDAMB-468 cells (E) by X-CELLigence RTCA DP technology (2-way ANOVA, ****P<0.0001, n=3). PBMCs were added at an effector to target ratio (E/T) of 5. (F) Schematic representation of Tras*anti-CD3-mediated tumor killing of human tumor organoids with endogenous T cells. (G) Bright field images of Tras*anti-CD3 induced killing of cancer cells in organoids from different patients. Scale bar: 50 μm. (H) Cell viability of the organoids from different patients (2-way ANOVA, ****P<0.0001, n=3). Data are shown as the means ± SDs. (I-J) The proportion of CD3+ T cells (I) and CD8+ T cells (J) of patient 1 after the addition of Tras*anti-CD3 (Student’s *t* test, *P<0.05, **P<0.01, n=3). (K) CD25 expression levels of CD8^+^ T cells of patient 1 after the addition of Tras*anti-CD3 (Student’s *t* test, *P<0.05, n=3).

Since Tras*anti-CD3 displayed desirable *in vitro* potency in 2D cell models, we tested whether the tissue penetration ability was influenced by the relatively large size of the Tras*anti-CD3 created by LacNAc-conju, which would hamper its clinical application with solid tumors^41^. It has been increasingly illustrated that patient-derived tumor models can predict potential clinical responses. Among them, tumor spheres or spheroids, as spherical aggregates from cancer cells, reflect more properties of solid tumors than that obtained with cells from 2D monolayer culture^42^. Therefore, we constructed patient-derived tumor-like cell clusters (PTCs)^43^, which is a model capable of recapitulating patients’ clinical responses with high accuracy, to evaluate the clinical potency of Tras*anti-CD3 (**Figure 6F**). In this study, PTCs generated from tumor single-cell suspensions of three bladder cancer patients were established, in which samples from patient 1 contained 6% T cells and showed HER2 expression on cancer cells (HER2 positive, CD3 high), patient 2 contained only 1% T cells and showed HER2 expression on cancer cells (HER2 positive, CD3 low), and patient 3 contained 0.3% T cells but showed no HER2 expression on cancer cells (HER2 negative, CD3 negative) (**Figure S11A-C**). As expected, Tras*anti-CD3 only exerted dose-dependent cellular cytotoxicity on PTCs from patient 1 (HER2 positive, CD3 high), and the cytotoxicity activity became saturated with 1.5 nM Tras*anti-CD3 (**Figure 6G-H**). In contrast, the tumor killing of PTCs from patient 2 (HER2 positive, CD3 low) was observed only with up to 60 nM Tras*anti-CD3 (****P<0.0001) (**Figure 6H, S11D**), suggesting that T cells were crucial for bispecific antibody-induced killing of solid tumors in clinical applications. For comparison, no tumor killing of PTCs was established with the samples (HER2 negative, CD3 negative) from patient 3 (**Figure 6G-H**). Furthermore, we used flow cytometry to analyze the phenotype of immune cell subtypes after the treatment of Tras*anti-CD3 in PTCs of patient 1. First, the frequency of CD3+ T cells inside PTCs increased from 31% to 54% after treatment (**Figure 6I, S11E**). In particular, the ratio of cytotoxic CD8+ T cells surged from 7% to 20% (**Figure 6J, S11F)**. Furthermore, the expression level of CD25 was significantly upregulated on CD8+ T cells (**Figure 6K, Figure S11G)**, indicating that Tras*anti-CD3 could activate T cells effectively, which further induced the release of granzyme B (GZMB) to initiate cancer cell death (**Figure S11H)**. Overall, the clinical potency of Tras*anti-CD3 toward PTC models confirmed that the Tras*anti-CD3 bispecific antibody constructed by LacNAc-conju could penetrate tissues to engage T cells and perform effective tumor killing; thus, this antibody exhibits great potential for clinical translation.

## Discussions and outlook

A robust and customized platform is necessary to perform site-specific modification of native IgG molecules in an attempt to meet the rapidly evolving applications. Glycoengineering is among the most convenient methods for performing site-specific and homogeneous construction of antibody conjugates from native IgG antibodies. However, with most of the current methods, hours to days are needed to complete the full conversion. The slow reaction rate is conventionally attributed to the unfavorable structure of donor substrate derivatives^44, 45^. Thus, engineering glycosyltransferases (GTs) and optimizing the chemical structure of the donor substrate are the main strategies to improve this type of approach.

In this work, we demonstrated that, despite being valuable, optimizing GT and donor substrate only improves unnatural fucosylation at a certain level. In contrast, reducing the complexity of Fc glycan, as the remodeling of acceptor substrates, could boost the reaction dramatically. More broadly, the monoantennary LacNAc structure could be introduced onto proteins beyond the antibody, such as at the C-terminus of the anti-PD-L1 nanobody, and then LacNAc could be recognized as a glycan tag by the FT enzyme. We demonstrated that the single LacNAc unit introduced onto the anti-PDL1 nanobody could be further modified by GDP-Fuc-AM-ssDNA with high efficiency **(Figure S12)**, suggesting that this glycan tag enables the enzymatic “click reaction”. Notably, LacNAc and GDP-Fuc as reaction handles of LacNAc-conju platform, compared to bioorthogonal handles, are much more hydrophilic. This unique property might help to accelerate the conjugation of hydrophobic and/or large payloads to antibodies, since the highly hydrophilic handles are easily exposed and help increase the payload solubility in aqueous buffer. Finally, the trisaccharide linker on the conjugate improved the pharmaceutical properties of the antibody conjugate, such as increasing the serum stability and reducing ADCC activity.

Taking advantage of the LacNAc-conju platform, we synthesized a bispecific antibody, Tras*anti-CD3, with a novel structure and demonstrated its excellent efficacy in patient-derived organoids. Our results support a recent finding that interdomain spacing and dual bivalency could improve bispecific antibody function^35^. The novel molecular structure of antibody conjugates from the LacNAc-conju platform provides great opportunities for developing antibody-biomacromolecule (X) conjugate (AXC). Ongoing efforts for further clinical translation of this technique focus on developing appropriate processing, such as enzyme fusions and immobilization. We expect that this platform will be widely used for constructing unlimited “AXC” drugs to provide therapeutics with novel structures and functions in the future.

## Methods

### Cell line culture and Mice

NCI-N87 cells, Jurkat Cells and NFAT-reporter-Jurkat cells were cultured in RPMI 1640 medium (Gibco) supplemented with 10% FBS (Gibco) and 1% penicillin/streptomycin. SKOV3, MDA-MB-231 cells and MDA-MB-468 cells were cultured in DMEM medium (Gibco) supplemented with 10% FBS (Gibco) and 1% penicillin/streptomycin (Gibco). SK-BR-3 cells were cultured in McCoy’s 5A medium (Gibco) supplemented with 10% FBS (Gibco) and 1% penicillin/streptomycin (Gibco). All cell lines were obtained from ATCC.

Human blood samples were collected from a healthy donor who provided informed consent on paper, and this study was approved by the Ethics Committee of Nanjing Drum Tower Hospital (2021-394-01).

BALB/c Nude mice were purchased from Hangzhou Medical College (Hangzhou). Both male and female mice of 4-5 weeks of age were used for most experiments. All purchased mice were specific pathogen-free (SPF) animals and were bred or housed under clean conditions. All animal studies were conducted in accordance with Institutional Animal Care and Use Committee guidelines and were performed at Hangzhou Medical College. (20210330Abbz0100018329)

### Human tumor samples

All human tissue samples were obtained from patients who provided informed consent on paper, and this study was approved by the Ethics Committee of Nanjing Drum Tower Hospital (2021-394-01).

### One-step synthesis chemoenzymatic synthesis of Tras-(A-LeX-Payload)_2_

Trastuzumab-(A-LacNAc)_2_ (3 mg/mL) was incubated with GDP-Fuc-AM-Payload (**2-6**) (80 μM) and α 1,3 FucT-2HR (0.8 mg/mL) in 25 mM Tris-HCl buffer (pH 7.5) with 20 mM MgCl_2_ at 30 °C for 1 hour. The modified trastuzumab was purified with protein A resin and confirmed by UPLC-TOF/MS analysis.

### PNGase F and PNGase A treatment

A solution of trastuzumab or modified trastuzumab (5 mg/mL) in PBS (pH 7.2) was incubated with PNGase F (0.1 mg/ml) and PNGase A (0.1 mg/ml) at 37 °C overnight.

All of the products were confirmed by UPLC-TOF/MS analysis.

### Release of N-linked glycans

A solution of trastuzumab or modified trastuzumab (5 mg/ml) in PBS (pH 7.2) was incubated with PNGase F (0.1 mg/ml) and PNGase A (0.1 mg/ml) at 37 °C overnight.

The reacted samples were precipitated with ice ethanol and centrifuged at 12,000 g, and the supernatant was aspirated. The supernatant was spin-dried with a speed vac and finally reconstituted with water for UPLC-TOF/MS analysis.

### ADCC activity assay

One days before the assay, the NFAT-reporter-Jurkat cells were seeded in RPMI 1640 medium and grown for one day. One day before the assay, SK-BR-3 cells at a density of 12,000 cells per well were seeded into a white clear-bottom 96-well microplate in 100 μL RPMI 1640 medium with a few empty wells (no cells) as the background luminescence control. On the day of the assay, the medium was removed from the plate, and antibodies were added with 60 μL RPMI 1640 medium (trastuzumab or modified trastuzumab). The cells were mixed with the antibodies at 37 °C in a CO_2_ incubator for 1 hour. NFAT-reporter-Jurkat cells were harvested by centrifugation and resuspended in RPMI 1640 medium at 75,000 Jurkat cells/40 μL. Each experiment was performed at least in triplicate. Then, 100 μl of RPMI 1640 medium was added to cell-free control wells to determine background luminescence. The plates were incubated at 37 °C in a CO_2_ incubator for 6 hours. Afterward, the luciferase assay was performed using the Bright-Lite Luciferase Assay System (Vazyme, DD1204-01) following the provided protocol. Finally, 100 μl Bright-Lite Luciferase reagent was added to each well and shaken gently at room temperature for ~30 minutes. The luminescence was measured using a luminometer. Data were analyzed by subtracting the average background luminescence (cell-free control wells) from the luminescence reading of all wells.

### One-step chemoenzymatic synthesis of Tras-(A-LeX-MMAE)_2_

Tras-(A-LacNAc)_2_ (3 mg/mL) was incubated with GDP-Fuc-AM-VcPAB-MMAE (80 μM) and α 1,3 FucT-2HR (0.8 mg/mL) in 25 mM Tris-HCl buffer (pH 7.5) with 20 mM MgCl_2_ at 30 °C for 1 hour. The modified trastuzumab was purified with protein A resin. All of the products were confirmed by UPLC-TOF/MS analysis.

### *In vitro* plasma stability assay of Tras-(A-LeX-MMAE)_2_

Human plasma was treated with protein A resin to remove the IgG. Then, the depleted IgG plasma was filter sterilized by a 0.22 μM filter. Tras-(A-LeX-MMAE)_2_ was incubated with the plasma to a final concentration of 100 μg/mL at 37 °C and 5% CO_2_. Samples were taken at 0, 2, 4, and 8 days and purified with protein A followed by UPLC-TOF/MS MS analysis.

### Synthesis of GDP-Fuc-AM-X nt ssDNA (X=20, 30, 42, 59, 88, 106)

Amine-modified ssDNA was produced by Sangon Co., Ltd, Shanghai. TCO-PEG_4_-NHS ester was purchased from Xi’an Confluore Biological Technology Co., Ltd. To make TCO-ssDNA, NH_2_-ssDNA and NHS-PEG4-TCO were reacted for 2 hours in 1 x borate buffered saline (BBS) solution (50 mM borate, 0.15 M NaCl, pH=8.5) containing 18% DMSO at final concentrations of 200 μM and 4 mM, respectively, and NHS-PEG4-TCO was added in equal amounts at half an hour intervals. The product TCO-ssDNA was then purified with DNA adsorption columns (Baiaolaibo Co., Ltd, Beijing). All of the products were confirmed by UPLC-TOF/MS analysis. GDP-Fuc-AM-MTz and TCO-ssDNA were reacted in 1:1 equivalent to obtain the final product GDP-Fuc-AM-ssDNA as this IEDDA reaction was completed quantitatively. The product TCO-ssDNA was then purified with DNA adsorption columns (Baiaolaibo Co., Ltd, Beijing). All of the products were confirmed by UPLC-TOF/MS analysis.

### One-step chemoenzymatic synthesis of Tras-(A-LeX-X nt ssDNA)2(x=20, 30, 42, 59, 88, 106)

Tras-(A-LacNAc)_2_ (1 mg/mL) was incubated with GDP-Fuc-AM-ssDNA (134 μM) and α 1,3 FucT-2HR (0.8 mg/mL) in 25 mM Tris-HCl buffer (pH 7.5) with 20 mM MgCl_2_ at 30 °C for 6 hours. The modified trastuzumab was purified with protein A resin. All of the products were confirmed by SDS‒ PAGE and UPLC-TOF/MS analysis.

### Synthesis of GDP-Fuc-AM-Anti-CD3 Nanobody (GF-Anti-CD3)

Anti-CD3 nanobody-LPETGG (1.5 mg/mL) was incubated with DBCO-PEG5-GGG (2 mM) and Sortase 5M ^39, 40^ (0.2 mg/mL) in PBS (pH 7.2) for 2 hours at r.t. The modified nanobody was purified with Ni-NTA resin. Anti-CD3 Nanobody-DBCO (10 mg/mL) was incubated with GDP-Fuc-AM-Az (666.7 μM) in 25 mM Tris-HCl buffer (pH 7.5) at 30 °C for 2 hours. The product GDP-Fuc-AM-Anti-CD3 Nanobody was then purified with a centrifugal filter unit (Millipore, 10 kDa MWCO, 0.5 mL sample volume). All of the products were confirmed by UPLC-TOF/MS analysis.

### One-step chemoenzymatic synthesis of Tras*anti-CD3

Tras-(A-LacNAc)_2_ (1 mg/mL) was incubated with GF-AntiCD3 (134 μM) and α 1,3 FucT-2HR (0.8 mg/mL) in 25 mM Tris-HCl buffer (pH 7.5) with 20 mM MgCl_2_ at 30 °C for 12 hours. The modified trastuzumab was purified with protein A resin. All of the products were confirmed by SDS‒PAGE and UPLC-TOF/MS analysis.

### Conjugate formation

Jurkat cells were incubated with 2.5 μM CFSE (Tonbo Biosciences, 13-0850-U500) (2.5 million/mL in PBS) for 10 minutes at room temperature, and SKOV3 cells were incubated with 2.5 μM eBioscience™ Cell Proliferation Dye eFluor™ 670 (Invitrogen, 65-0840-85) (2.5 million/ml in PBS) for 10 minutes at room temperature. After washing three times with FACS buffer (DPBS containing 2% FBS), Jurkat cells and SKOV3 cells were mixed (20000:20000) and added to serially titrate Tras*anti-CD3 in duplicate. After 15 minutes of incubation at 37 °C, flow cytometry analysis was performed directly using an Agilent Novocyte Quanton flow cytometer, and analysis was performed using FlowJo 10.5.3.

### Activation assay

Human blood samples were collected from a healthy donor who provided informed consent on paper, and this study was approved by the Ethics Committee of Nanjing Drum Tower Hospital (2021-394-01). Human peripheral blood mononuclear cells (PBMCs) were obtained by Ficoll (Ficoll-Paque Plus, GE) density gradient centrifugation. PBMCs (100,000) were incubated with SKOV3 cells (20,000) in 96-well flat-bottom plates, in which 15 nM Tras*anti-CD3 bispecific antibody or control Anti-CD3 Nanobody was added. After 48 hours, the supernatant was collected and frozen at −80 °C for subsequent analysis. Cell pellets were then stained with antibodies against CD4-PB (Biolegend, Clone OKT4, 317424), CD8a-PE (Biolegend, Clone HIT8a, 300908), CD3-Alexa Fluor® 700 (Biolegend, Clone OKT3, 317340), CD69-APC (Biolegend, Clone FN50, 310910), and CD25-Percp/Cy5.5 (Biolegend, Clone BC96, 302626) to assess CD69 and CD25 upregulation. Cells were acquired using an Agilent Novocyte Quanton and analyzed using FlowJo 10.5.3. Graphs were obtained using GraphPad Prism 8.

### Cytokine measurements

Frozen supernatant from the activation assay (48 hours) was used to measure *in vitro* cytokine production. Briefly, the supernatant was diluted 5-fold. The ELISA kits for human IFN-γ (4A Biotech, CHE0017) and TNF-α (4A Biotech, CHE0019) were taken from the refrigerator 30 minutes before use. Various solutions were prepared according to the preparation work. The double-sandwich method was used for testing. The samples were incubated separately, and biotin and enzyme conjugates were added. Subsequently, chromogenic reagents were added and incubated in the dark for 15 minutes. Finally, stop solution was added to test the OD450 value immediately. Data were analyzed using GraphPad Prism 8.

### Cytotoxicity measurements of target cells by effector cells

Cytotoxicity measurements of target cells by effector cells were tested using the RTCA-DP instrument. Then, 50 μL of DMEM was added to each well of an E-Plate 16 PET plate and put into an RTCA machine to measure the baseline, and the CI value was within ±0.063. Then, SKOV3 cells (10,000) or MDAMB468 cells (20,000) in each well were added. When the cells grew to the late logarithmic growth phase, the plate was removed, and PBMCs (50,000) and 15 nM Tras*anti-CD3 were added and monitored. Each experiment was performed in duplicate.

### Culture of PTCs from solid tumors

Collected samples were processed in ice-cold PBS using methods described in the literature^43^. Treated cells were cultured in a 37 °C, 5% CO_2_ incubator. The PTC growth medium with 100 IU IL2 was renewed every 3 days as necessary. This study was approved by the Ethics Committee of Nanjing Drum Tower Hospital (2021-394-01).

### Cytotoxicity measurements of PTCs

PTCs were collected, centrifuged at 300 × g and 4 °C for 10 minutes, washed with PBS, and then resuspended in G/I-PTC growth medium. Then, 100 μL of medium containing 50-100 PTCs was inoculated into the supernatant in a 96-well plate with low adsorption. Next, 50 μL of PTC growth medium containing dual antibodies was added to the wells at different drug concentrations. The plates were incubated at 37 °C and 5% CO_2_. After 5 d of drug treatment, cell viability was detected with a Cell-Counting-Lite® 3D Luminescent Cell Viability Assay (Vazyme).

### Activation assay of tumor-infiltrating T cells in PTC

Following the steps above, 100 μL of medium containing 50-100 PTCs was inoculated with 60 nM Tras*anti-CD3 in a 37 °C, 5% CO_2_ incubator for 5 days. Then, the PTCs were digested using TrypLe (Thermo Fisher Scientific). Briefly, the PTCs were dissociated into single cells by pipetting every 2 minutes. After they were washed with DMEM (Thermo Fisher Scientific), the cells were centrifuged at 300 × g and 4 °C for 10 minutes. Cell pellets were then stained with antibodies against CD4-PB (Biolegend, Clone OKT4, 317424), CD8a-PE (Biolegend, Clone HIT8a, 300908), CD3-Alexa Fluor® 700 (Biolegend, Clone OKT3, 317340), CD69-APC (Biolegend, Clone FN50, 310910), and CD25-Percp/Cy5.5 (Biolegend, Clone BC96, 302626) to assess CD69 and CD25 upregulation. Cells were acquired using an Agilent Novocyte Quanton and analyzed using FlowJo 10.5.3. Graphs were obtained using GraphPad Prism 8. This study was approved by the Ethics Committee of Nanjing Drum Tower Hospital (2021-394-01).

## Acknowledgments

J.P.L. acknowledged the support from the National Key R&D Program of China (2019YFA09006600), National Natural Science Foundation of China (21977048, 92053111), the Fundamental Research Funds for the Central Universities, Natural Science Foundation of Jiangsu Province (BK20202004), Beijing National Laboratory for Molecular Sciences (BNLMS202008), Program for Innovative Talents and Entrepreneur in Jiangsu, and the Excellent Research Program of Nanjing University (ZYJH004). We thank Prof. Jianzhong Xi for his kind guidance in PTC construction.

## Author contributions

J.P.L., Y.Y., Y.Y., and Z.S. designed the experimental strategies. Y.Y., Z.S., Z.Z., J.C., and J.H.

performed the experiments with the help of J.C., J.H., X.J., G.Y., Q.X., X.Z., and W.S., Y.Y., T.T., J.P.L., and Y.Y. prepared the figures. J.P.L., Y.Y., Y.Y., and T.T. wrote the manuscript. All authors analyzed the data and edited the manuscript.

## Additional information

Correspondence and requests for materials should be addressed to J.P. L or Y.Y.

## References

1. Beck, A., Goetsch, L., Dumontet, C. & Corvaia, N. Strategies and challenges for the next generation of antibody-drug conjugates. Nat Rev Drug Discov 16, 315–337 (2017).

2. Modi, S. et al. Trastuzumab deruxtecan in previously treated HER2-low advanced breast cancer. N Engl J Med 387, 9–20 (2022).

3. Mullard, A. Antibody-oligonucleotide conjugates enter the clinic. Nat Rev Drug Discov 21, 6–8 (2022).

4. Stoeckius, M. et al. Simultaneous epitope and transcriptome measurement in single cells. Nat Methods 14, 865–868 (2017).

5. Robinson, P.V., Tsai, C.T., de Groot, A.E., McKechnie, J.L. & Bertozzi, C.R. Glyco-seek: ultrasensitive detection of protein-specific glycosylation by proximity ligation polymerase chain reaction. J Am Chem Soc 138, 10722–10725 (2016).

6. Bapat, S.P. et al. Obesity alters pathology and treatment response in inflammatory disease. Nature 604, 337–342 (2022).

7. Hudak, J.E. et al. Synthesis of heterobifunctional protein fusions using copper-free click chemistry and the aldehyde tag. Angew Chem Int Ed Engl 51, 4161–4165 (2012).

8. Xiao, H., Woods, E.C., Vukojicic, P. & Bertozzi, C.R. Precision glycocalyx editing as a strategy for cancer immunotherapy. Proc Natl Acad Sci USA 113, 10304–10309 (2016).

9. Gray, M.A. et al. Targeted glycan degradation potentiates the anticancer immune response *in vivo*. Nat Chem Biol 16, 1376–1384 (2020).

10. Agarwal, P. & Bertozzi, C.R. Site-specific antibody-drug conjugates: the nexus of bioorthogonal chemistry, protein engineering, and drug development. Bioconjug Chem 26, 176–192 (2015).

11. Lu, H. et al. Site-specific antibody-polymer conjugates for siRNA delivery. J Am Chem Soc 135, 13885–13891 (2013).

12. Deshaies, R.J. Multispecific drugs herald a new era of biopharmaceutical innovation. Nature 580, 329–338 (2020).

13. Wacker, C., Berger, C.N., Girard, P. & Meier, R. Glycosylation profiles of therapeutic antibody pharmaceuticals. Eur J Pharm Biopharm 79, 503–507 (2011).

14. Chudasama, V., Maruani, A. & Caddick, S. Recent advances in the construction of antibody-drug conjugates. Nat Chem 8, 114–119 (2016).

15. van Geel, R. et al. Chemoenzymatic conjugation of toxic payloads to the globally conserved N-glycan of native mabs provides homogeneous and highly efficacious antibody-drug conjugates. Bioconjug Chem 26, 2233–2242 (2015).

16. Zeglis, B.M. et al. Enzyme-mediated methodology for the site-specific radiolabeling of antibodies based on catalyst-free click chemistry. Bioconjug Chem 24, 1057–1067 (2013).

17. Li, X., Fang, T. & Boons, G.J. Preparation of well-defined antibody-drug conjugates through glycan remodeling and strain-promoted azide-alkyne cycloadditions. Angew Chem Int Ed Engl 53, 7179–7182 (2014).

18. Huang, W., Giddens, J., Fan, S.Q., Toonstra, C. & Wang, L.X. Chemoenzymatic glycoengineering of intact IgG antibodies for gain of functions. J Am Chem Soc 134, 12308–12318 (2012).

19. Tang, F. et al. One-pot N-glycosylation remodeling of IgG with non-natural sialylglycopeptides enables glycosite-specific and dual-payload antibody-drug conjugates. Org Biomol Chem 14, 9501–9518 (2016).

20. Parsons, T.B. et al. Optimal synthetic glycosylation of a therapeutic antibody. Angew Chem Int Ed Engl 55, 2361–2367 (2016).

21. Li, C. et al. Site-selective chemoenzymatic modification on the core fucose of an antibody enhances its Fcγ receptor affinity and ADCC activity. J Am Chem Soc 143, 7828–7838 (2021).

22. Shi, W. et al. One-step synthesis of site-specific antibody-drug conjugates by reprograming IgG glycoengineering with LacNAc-based substrates. Acta Pharm Sin B 12, 2417–2428 (2022).

23. Zhang, X. et al. Site-specific chemoenzymatic conjugation of high-affinity M6P glycan ligands to antibodies for targeted protein degradation. ACS Chem Biol DOI: 10.1021/acschembio.1c00751 (2022).

24. Zhang, X. et al. General and robust chemoenzymatic method for glycan-mediated site-specific labeling and conjugation of antibodies: facile synthesis of homogeneous antibody-drug conjugates. ACS Chem Biol 16, 2502–2514 (2021).

25. Li, J. et al. A single-step chemoenzymatic reaction for the construction of antibody-cell conjugates. ACS Cent Sci 4, 1633–1641 (2018).

26. Sun, H.Y. et al. Structure and mechanism of Helicobacter pylori fucosyltransferase. A basis for lipopolysaccharide variation and inhibitor design. J Biol Chem 282, 9973–9982 (2007).

27. Lin, Y.-N. et al. Chemoenzymatic synthesis of GDP-L-Fucose derivatives as potent and selective α-1,3-fucosyltransferase inhibitors. Adv Syn Catal 354, 1750–1758 (2012).

28. Cobb, B.A. The history of IgG glycosylation and where we are now. Glycobiology 30, 202–213 (2020).

29. Trastoy, B. et al. Crystal structure of Streptococcus pyogenes EndoS, an immunomodulatory endoglycosidase specific for human IgG antibodies. Proc Natl Acad Sci USA 111, 6714–6719 (2014).

30. Klontz, E.H. et al. Structure and dynamics of an α-fucosidase reveal a mechanism for highly efficient IgG transfucosylation. Nat Commun 11, 6204 (2020).

31. Lyon, R.P. et al. Reducing hydrophobicity of homogeneous antibody-drug conjugates improves pharmacokinetics and therapeutic index. Nat Biotechnol 33, 733–735 (2015).

32. Blackman, M.L., Royzen, M. & Fox, J.M. Tetrazine ligation: fast bioconjugation based on inverse-electron-demand Diels-Alder reactivity. J Am Chem Soc 130, 13518–13519 (2008).

33. Labrijn, A.F., Janmaat, M.L., Reichert, J.M. & Parren, P. Bispecific antibodies: a mechanistic review of the pipeline. Nat Rev Drug Discov 18, 585–608 (2019).

34. Brinkmann, U. & Kontermann, R.E. Bispecific antibodies. Science 372, 916–917 (2021).

35. Santich, B.H. et al. Interdomain spacing and spatial configuration drive the potency of IgG-[L]-scFv T cell bispecific antibodies. Sci Transl Med 12 (2020).

36. Taylor, R.J. et al. π-clamp-mediated homo- and heterodimerization of single-domain antibodies via site-specific homobifunctional conjugation. J Am Chem Soc 144, 13026–13031 (2022).

37. Taylor, R.J., Geeson, M.B., Journeaux, T. & Bernardes, G.J.L. Chemical and enzymatic methods for post-translational protein-protein conjugation. J Am Chem Soc 144, 14404–14419 (2022).

38. Yuan, D. et al. Site-selective lysine acetylation of human immunoglobulin G for immunoliposomes and bispecific antibody complexes. J Am Chem Soc DOI: 10.1021/jacs.2c07594 (2022).

39. Ge, Y. et al. Enzyme-mediated intercellular proximity labeling for detecting cell-cell interactions. J Am Chem Soc 141, 1833–1837 (2019).

40. Chen, L. et al. Improved variants of SrtA for site-specific conjugation on antibodies and proteins with high efficiency. Sci Rep 6, 31899 (2016).

41. Minchinton, A.I. & Tannock, I.F. Drug penetration in solid tumours. Nat Rev Cancer 6, 583–592 (2006).

42. Costa, E.C. et al. 3D tumor spheroids: an overview on the tools and techniques used for their analysis. Biotechnol Adv 34, 1427–1441 (2016).

43. Yin, S.Y. et al. Patient-derived tumor-like cell clusters for drug testing in cancer therapy. Sci Transl Med 12 (2020).

44. Choi, J. et al. Engineering orthogonal polypeptide GalNAc-transferase and UDP-sugar pairs. J Am Chem Soc 141, 13442–13453 (2019).

45. Tian, Y. et al. One-step enzymatic labeling reveals a critical role of O-GlcNAcylation in cell-cycle progression and dna damage response. Angew Chem Int Ed Engl 60, 26128–26135 (2021).

